# SARS-CoV-2 specific memory T lymphocytes from COVID-19 convalescent donors: identification, biobanking and large-scale production for Adoptive Cell Therapy

**DOI:** 10.1101/2020.10.23.352294

**Authors:** C Ferreras, B Pascual-Miguel, C Mestre-Durán, A Navarro-Zapata, L Clares-Villa, C Martín-Cortázar, R De Paz, A Marcos, JL Vicario, A Balas, F García-Sánchez, C Eguizabal, C Solano, M Mora-Rillo, B Soria, A Pérez-Martínez

**Affiliations:** Hospital La Paz Institute for Health Research, IdiPAZ, La Paz University Hospital, Madrid, Spain; University Hospital La Paz, Hematology Department, Madrid, Spain; Histocompatibility. Centro de Transfusión de Madrid. Madrid, Spain; Research Unit, Basque Center for Blood Transfusion and Human Tissues, Osakidetza, Galdakao, Bizkaia, Spain; Cell Therapy, Stem Cells and Tissues Group, Biocruces Bizkaia Health Research Institute, Barakaldo, Bizkaia, Spain; Hospital Clínico Universitario de Valencia/Instituto de Investigación Sanitaria INCLIVA, Universidad de Valencia; Infectious Diseases Unit, Internal Medicine Department, University Hospital La Paz, - Hospital La Paz Institute for Health Research, IdiPAZ, La Paz University Hospital, Madrid, Spain; Instituto de Bioingeniería, Universidad Miguel Hernández de Elche, Alicante; Instituto de Investigación Sanitaria Hospital General y Universitario de Alicante ISABIAL; University Hospital La Paz, Pediatric Hemato-oncology Department, Madrid, Spain; Faculty of Medicine Universidad Autónoma de Madrid, Madrid, Spain

## Abstract

SARS-CoV-2 is causing a second outbreak so the hope for its complete eradication is far from happening. In the absence of effective vaccines, it is mandatory to find effective treatments with low adverse effects able to treat hospitalized patients with COVID-19 disease. In this work, we determined the existence of SARS-CoV-2 specific T cells within the CD45RA^−^ T memory cells from the blood of convalescent donors. Memory T cells can respond quickly to the infection and provide long-term immune protection to reduce the severity of the COVID-19 symptoms. Also, CD45RA^−^ memory T cells confer protection from other pathogens the donors encountered in their life. This is vital to clear other secondary infections usually developed in hospitalized COVID-19 patients. SARS-CoV-2 specific memory T cells were found within all the CD45RA^−^ subsets CD3^+^, CD4^+^, CD8^+^, and in the central memory and effector memory subpopulations. The procedure to obtain the cells is feasible, easy to implement for small scale manufacture, quick and cost-effective involving minimal manipulation, and without GMP condition requirements. This biobank of specific SARS-CoV-2 memory T cells would be immediately available ‘off-the-shelf’ to treat moderate/severe cases of COVID-19 increasing the therapeutic options available for these patients.

## INTRODUCTION

The new severe acute respiratory syndrome coronavirus 2, or SARS-CoV-2 has emerged causing a worldwide pandemic from late 2019. This coronavirus is causing an infectious disease called COVID-19 with a wide range and diverse symptoms. In most infected patients the illness generates mild symptoms including fever and cough but, in some others, causes a life-threatening disease with symptoms that include pneumonia, dyspnea and a hyperinflammatory process which includes cytokine storm, systemic immunethrombosis, etc. Patients suffering from these symptoms need hospitalization and treatment. A common feature of this severe disease is lymphopenia making them more vulnerable to co-infections and correlating with the severity of the disease^1,2^. With strong restrictive measures, social distancing, and healthcare interventions the first wave was contained, although thousands of patients died. Far from disappearing SARS-CoV-2 is causing a second wave thus diminishing the hope of its complete eradication. Vaccine development actively pursues to generate active immunity through vaccine immunization^3^ but there is uncertainty about how long antibody-mediated immune response to COVID-19 will last^4^. We need effective treatments that can reduce the severity of symptoms, time of hospitalization and increase survival.

So far, the only treatment for COVID-19 is supportive. Antiviral treatment with lopinavir–ritonavir is not effective in improving outcomes for hospitalized patients with COVID-19^5^; also, remdesivir has been recently approved to treat COVID-19 although its beneficial effect is still controversial^6,7^. Preliminary results with anti-inflammatory therapies such as dexamethasone^8^ or mesenchymal stromal cells^9^ have shown promising results for critical patients (WHO Grade 6 and 7)^10^, but we do not have yet an effective antiviral therapy able to stop the progress of this diseases at early stages: (WHO Grade 1-4 moderate and severe) or even to prevent COVID-19.

The role of the adaptive immunity in COVID-19 disease and the protective immunity conferred by T cells is being characterized^11–15^ but the role of memory T cells conferring protection against SARS-CoV-2 has not been properly defined yet. The existence of memory T cells specific for another SARS coronavirus was found up to 11 years post-infection^16^. This immunological memory creates a more rapid and robust secondary immune response to reinfections. This fact is determinant and constitutes the basis of adoptive cell therapy of viral infections in immunosuppressed patients in the context of allogeneic hematopoietic stem cell transplantation (HSCT). In HSCT the infusion of CD45RA^−^ memory T cells has considerably reduced the morbidity and mortality induced by viral reactivations such as cytomegalovirus (CMV) or Epstein Barr Virus (EBV) and, at the same time, has reduced the alloreactivity conferred by the naïve CD45RA^+^ T cells^17–21^.

Memory T cells do appear when T lymphocytes recognize a pathogen presented by their local antigen-presenting cells. These T cells activate, proliferate, and differentiate into effector cells secreting molecules to control the infection. Once the pathogen has been cleared most of the antigen-specific T cells disappear and a pool of heterogeneous long-lived memory T cells persist^22,23^. This population of memory T cells defined as CD45RA^−^ or CD45RO^+^ is maintained over time conferring a quick and long-term immune protection against subsequent reinfections^24,25^.

In this work, we report the existence of a SARS-CoV-2 specific T cell population within the CD45RA^−^ T memory cells from the blood of convalescent donors that can be easily, effectively, and rapidly isolated by CD45RA depletion. These specific SARS-CoV-2 CD45RA^−^ T memory cells may be able to clear virally infected cells and to confer T cell immunity for subsequent reinfections. These cells can be stored to be used in moderate and severe cases of hospitalized COVID-19 patients becoming an off-the-self living drug.

## METHODS

### Donors’ characteristics

**Table 1** shows the donors’ characteristics. Six COVID-19 convalescent donors and two healthy controls were included. The convalescent donors were all tested for SARS-CoV-2 by reverse-transcriptase polymerase chain reaction (RT-PCR) in nasopharyngeal samples between March and April 2020. Eligibility criteria included age 21 to 65, a history of COVID-19 with documented positive RT-PCR test for SARS-CoV-2. At the time of this study, they were all tested negative for SARS-CoV-2. The median age of the convalescent donors was 37 years old (range 23-41), three were females and three were males. The median days until PCR negative for SARS-CoV-2 was 13 days (range 5-17). Two of them presented with bilateral pneumonia but no hospitalization was needed. Just one of them received treatment with oral hydroxycloroquine + azytromicine + lopinariv/ritonavir and another one was treated with oral hydroxycloroquine + azytromicine. Two healthy donors were enrolled who have not been exposed to COVID-19 patients and were tested negative for anti-SARS-CoV-2 antibodies in June 2020. All the participants gave written consent with approval by the Hospital Institution Review Board (IRB number: 254/20)

**Table 1.** Participants’ characteristics.

|  |  |
| --- | --- |
| Number of donors | 6 |
| Mean age (range) | 37 (23-41) |
| Gender (female/male) | 3/3 |
| Days to SARS-CoV-2 negative PCR (range) | 12.83 (5-17) |
| Bilateral pneumonia, n(%) | 2 (33.3%) |
| Days until samples taken from negative PCR (range) | 14.70 (7-33) |
| Ambulatory, n (%) | 6 (100%) |
| **Treatment** |  |
| Acetaminophen | 4 |
| Hydroxychloroquine+azytromicine+lopinariv/ritonavir | 1 |
| Hydroxychloroquine+azytromicine | 1 |

### Cell processing and detection of SASR-CoV-2 specific memory T cells by IFN-γ assay

Peripheral Blood Mononuclear Cells (PBMCs) from healthy donors and the convalescent donors were isolated from their peripheral blood by density gradient centrifugation using Ficoll-Paque (GE Healthcare, Illinois, Chicago, USA). Briefly, cells were rested overnight (o/n) at 37°C in TexMACS Medium (Miltenyi Biotec, Bergisch Gladbach, Germany) supplemented with 10% AB serum (Sigma-Aldrich, Saint Louis, Missouri, USA) and 1% Penicillin/Streptomycin (Sigma-Aldrich, Saint Louis, Missouri, USA). The next day 1×10^6^ cells were stimulated with pooled or individual overlapping SARS-CoV-2 peptides at a final concentration of 0.6 nmol/ ml. For positive control, 1×10^6^ cells were stimulated in the presence of plate-bound stimulator OKT3 at a final concentration of 2.8 μg/ml (mouse anti-human CD3 Clone OKT3 BD Biosciences). Cells with SARS-CoV-2 peptides and positive control were co-stimulated with CD28/CD49d at a final concentration of 5 μg/ml (anti-human CD28/CD49d Purified Clone L293 L25 BD Biosciences). Basal IFN-γ production by PBMCs was included as background control in the absence of stimulation and co-stimulation. The peptide pools were short 15-mer peptides with 11 amino acid overlaps which can bind MHC class I and class II complexes and thus were able to stimulate both CD4^+^ and CD8^+^ T cells. The peptides cover the immunodominant sequence domains of the surface glycoprotein S, and the complete sequence of the nucleocapsid phosphoprotein N and the membrane glycoprotein M (GenBank MN908947.3, Protein QHD43416.1, Protein QHD43423.2, Protein QHD43419.1) (Miltenyi Biotec, Germany). After 5 hours of stimulation, the cells were labeled with the IFN-γ Catch Reagent (IFN-γ Secretion Assay-Detection Kit, human Miltenyi Biotec) containing bispecific antibodies for CD45 and IFN-γ which was secreted by the stimulated target cells. After the secretion phase, the cell surface-bound IFN-γ was targeted using the IFN-γ PE antibody included in the kit.

### Phenotype of memory T cells containing SARS-CoV-2 specific T cells by flow cytometry assay

Analysis of the cell composition in the T cells specifically activated by SARS-Cov-2 in the IFN-γ assay was performed by subtracting the basal cytokine response from the background control. Cell surface staining was performed for 20 minutes at 4 degrees with the following fluorochrome-conjugated antibodies titrated to their optimal concentrations:, CD45RA FITC (BD Pharmingen), CD27 APC (BD Pharmingen), CD3 viogreen (Miltenyi Biotec), CD4 PECy7 (BD Pharmingen), CD8 APC Cy7 (BD Pharmingen), 7AAD (BD Horizon). For the Treg panel CD25 BV421 (BD Horizon), and CD127 PE-CF594 (BD Horizon), were used. For the activation panel HLA-DR BV 421 (BD Pharmingen), CD69 BV421 (Biolegend), CD25 BV421 (BD Horizon), were used. For the exhaustion panel PD1 AF700 (Biolegend), NKG2A BV421 (Biolegend) were used. For the chemokine panel, CD103 BV421 (BD Horizon) and CCR7 PE-CF594 (BD Horizon) were used. An average of 200.000 cells was acquired. Cell acquisition was performed using a Navios cytometer (Beckman Coulter). The analysis was performed using FlowJo 10.7.1 (FlowJo LLC).

### IL15 stimulation of memory T cells

CD45RA^−^ memory T lymphocytes from the convalescent donor were thawed and stimulated with IL15 to obtain an activated phenotype. Cells were incubated in TexMACS Medium (Miltenyi Biotec, Germany) supplemented with 5% AB serum (Sigma-Aldrich, Saint Louis, Missouri, USA) and 1% Penicillin/Streptomycin (Sigma-Aldrich, Saint Louis, Missouri, USA) plus 50 ng/ml of IL15 o/n and for 72 hours. After that time the cells were harvested and the phenotypic assay was performed. The same culture without IL15 was run in parallel as a control.

### Donor selection, HLA typing, and large clinical scale CD45RA^+^

T cell depletion Criteria for the convalescent donor selection were: secretion of IFN-γ upon activation with the 3 SARS-CoV-2 specific peptides (M, N, S) and the most frequent HLA typing to cover most of the population. The HLA phenotype of the convalescent donor was done at the Centro de Transfusión of the Comunidad of Madrid on two independent samples by SSO and NGS: A*02:01,A*24:02 / B*44:02,B*51:01 / C*16:02,C*16:04 / DRB1*07:01,DRB1*11:03 / DQB1*02:02,DQB1*03:01.

Non-mobilized apheresis was performed at the Bone Marrow Transplant and Cell Therapy Unit for University Hospital La Paz, Madrid, Spain by using CliniMACS Plus device (Milteny Biotec). The donor gave written informed consent by the Declaration of Helsinki protocol, and the study was performed according to the guidelines of the local ethics committee (IRB Number 5579). The donor complied with the requirements regarding quality and safety for the donation, obtaining, storage, distribution, and preservation of human cells and tissues under the Spanish specific regulation. Following apheresis, CD45RA^+^ cells were depleted by immunomagnetic separation using a CliniMACS CD45RA Reagent and CliniMACS Plus system, both from Miltenyi Biotec, following manufacturer instructions. CD45RA^−^ cells were frozen using autologous plasma plus 5% dimethyl sulfoxide (DMSO) and stored. We were able to cryopreserve 30 aliquots at different doses according to the trial design. The viability, purity, phenotype and spectratyping of CD45RA^−^ fraction were analyzed by flow cytometry (FCM).

### TCR spectratyping

Most of the CDR3-encoding regions of the TCRV-β and -γ genes were amplified using 2 V-J multi-primer sets for each locus and one additional multi-primer set covering D-J TCRV-β (Vitro, Master Diagnostica, Spain). Primers marked at their 5’ end with 6-FAM fluorochrome, allowed denatured fragment size analysis by capillary electrophoresis (ABI3130 DNA-analyzer) and Genemapper software (Thermo Fisher Scientific, USA).

### Statistical analysis

Quantitative variables are expressed as Mean + Standard Deviation (SD). Qualitative variables are expressed in percentages (%). Two-tailed Mann-Whitney non-parametric test was used for comparison means for non-paired samples using GraphPad Prism (version 8.0.0 for Windows, GraphPad Software, San Diego, California USA). A P value < 0.05 was considered statistically significant.

## RESULTS

### Memory T cells from convalescent donors contain a SARS-CoV-2 specific population

PBMCs of the convalescent and healthy donors were stimulated with the three overlapping peptides (M, N, S) specific of SARS-CoV-2 for 5 hours. We detected the existence of SARS-CoV-2 specific population by the production of IFN-γ in both subsets naïve CD45RA^+^ and memory CD45RA^−^ T cells in the PBMCs of the convalescent donors but not in healthy controls (**Table 2** and **Figure 1**). They also showed reactivity for the single peptides M, N, and S (data not shown). The mean CD45RA^−^CD3^+^ population in the convalescent donors was close to 90%. Expression of IFN-γ within CD45RA^−^CD3^+^ population was 1.12% whereas the expression of IFN-γ within CD45RA^+^CD3^+^ population was 0.40% (P=0.065) (Table 2). We didn’t detect the expression of IFN-γ in healthy individuals. Despite is a small cohort we didn’t find a synergistic effect on the percentage of IFN-γ when the three peptides are mixed when compared to the single peptides (data not shown).

**Table 2.**
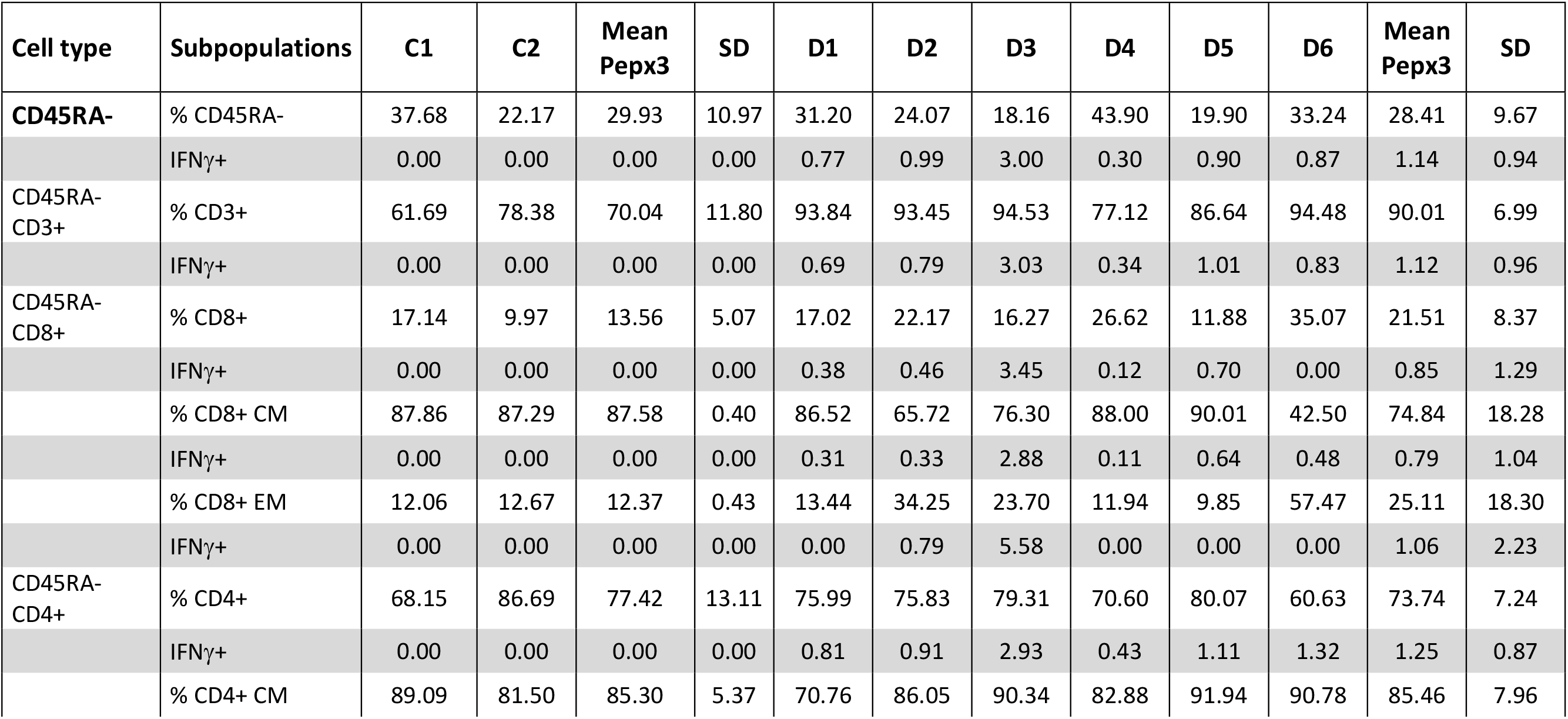
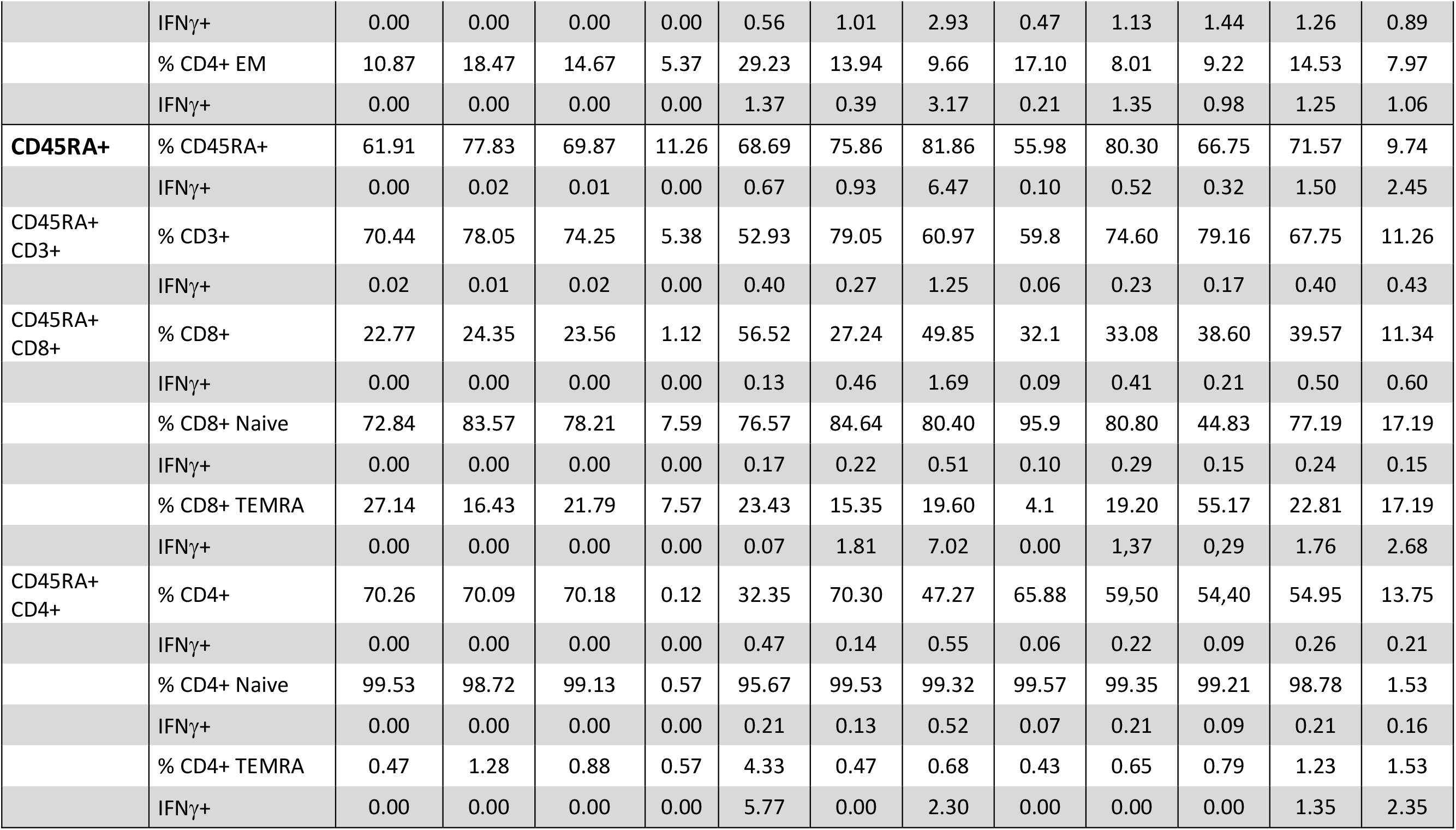
Immunophenotypic characterization of the healthy controls and the convalescent donors after co-culture with the mixture of the three SARS-CoV-2 peptides (M, N, S)

**Figure 1.**
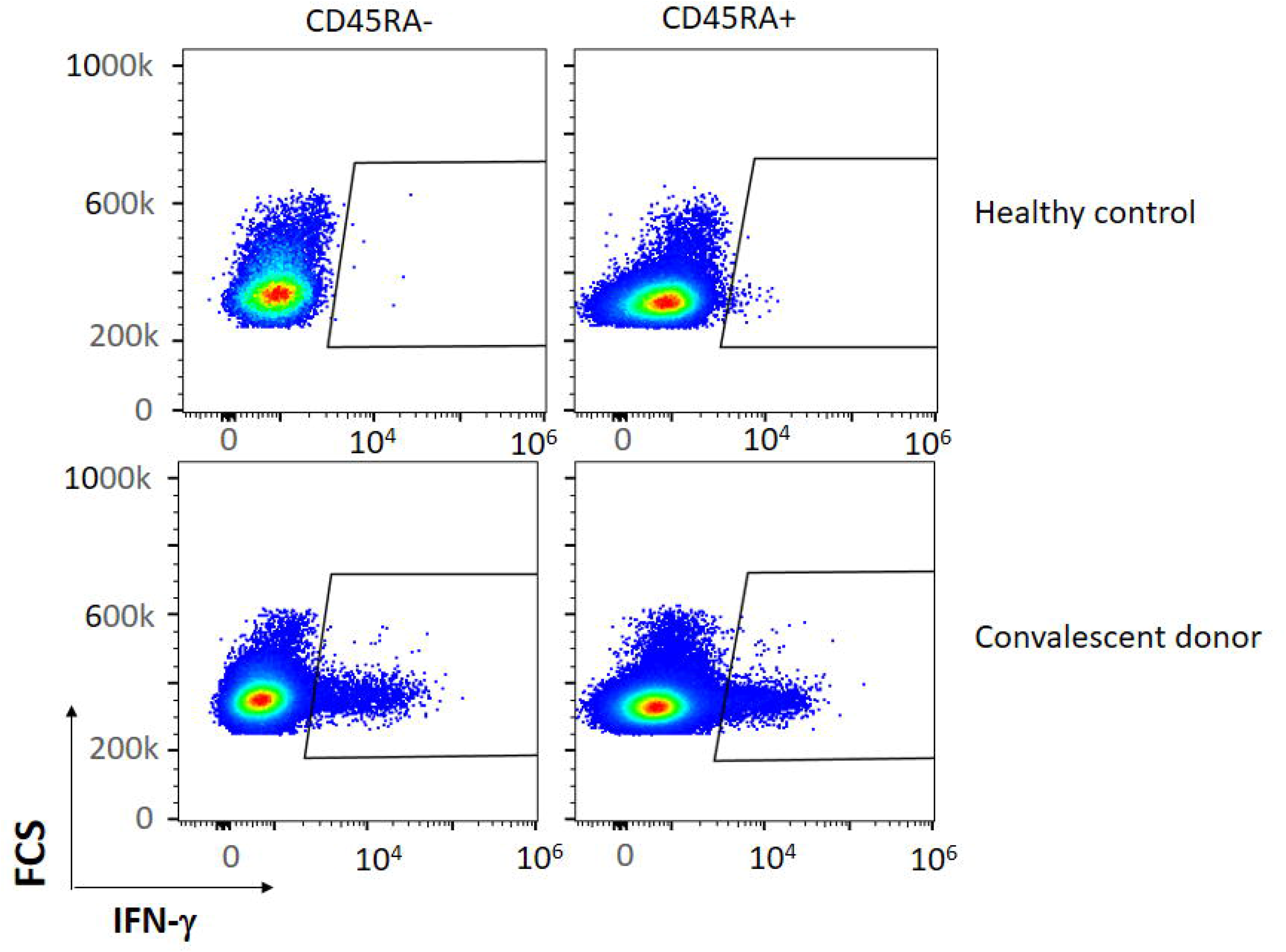
Representative figure of the expression of IFN-γ in a convalescent and healthy donor within the CD45RA^+^ and CD45RA^−^ subpopulations after co-culture with the mixture of the three peptides (M, N, S)

### Identification of SARS-CoV-2 specific CD4^+^ T and CD8^+^ T cell responses within CD45RA-memory T cells

Then, we asked whether both CD8^+^ and CD4^+^ subsets contained specific SARS-CoV-2 T cells within the PBMCs of the convalescent donors. Among all subsets studied we observed that CD45RA-CD4^+^ and CD45RA-CD8^+^ cells expressed 1.25% and 0.85% of IFN-γ respectively (P= 0.132). (Table 2, Figure 2, and data not shown). Thus, all the convalescent donors recovered from COVID-19 generated CD4^+^ T and CD8^+^ T cells responses against SARS-CoV-2 within the memory CD45RA-T cell population.

**Figure 2.**
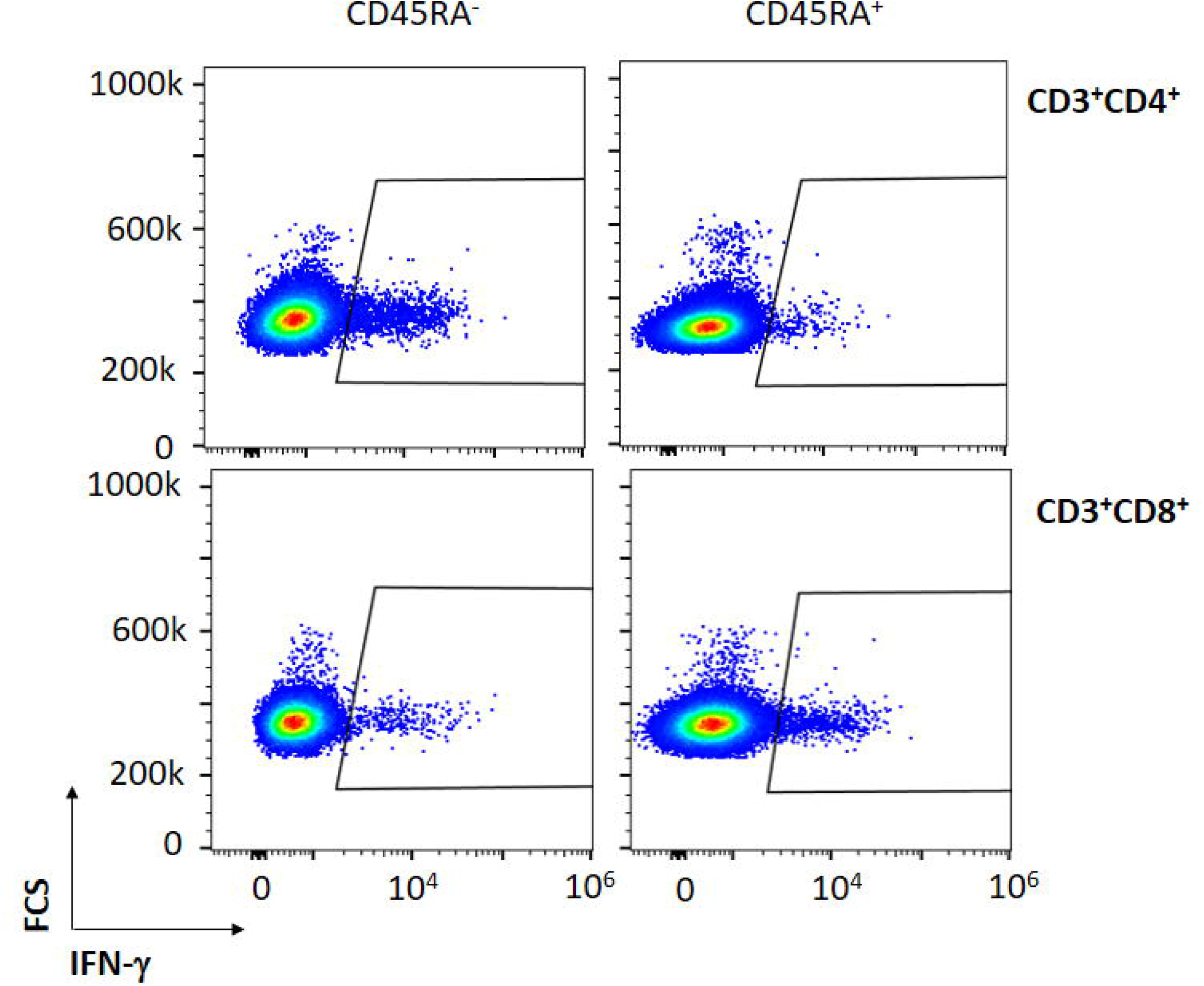
Representative figure of the expression of IFN-γ in a convalescent donor within the CD45RA^−^CD3^+^CD4^+^ and CD45RA^−^CD3^+^CD8^+^ subpopulations after co-culture with the mixture of the three peptides (M, N, S)

We then analysed the T central memory (CM) (CD45RA^−^CD3^+^CD27^+^) and T effector memory (EM) (CD45RA^−^CD3^+^CD27^−^) compartments. While no significant differences were detected, we observed responses to the SARS-CoV-2 specific peptides within all subpopulations. Within the CD4^+^ and CD8^+^ CM T cell subsets we detected a mean of 1.26% and 0.79% of IFN-γ^+^ cells respectively (P=0.132). Looking at the CD4^+^ and CD8^+^ EM T cell subsets we detected a mean of 1.25% and 1.06% of IFN-γ^+^ cells respectively (P=0.108). (**Table 2** **and** **Figure 3**). These data show the existence of a population of memory T cells specific for SARS-CoV-2 within the CD45RA^−^CD3^+^ memory T cells.

**Figure 3.**
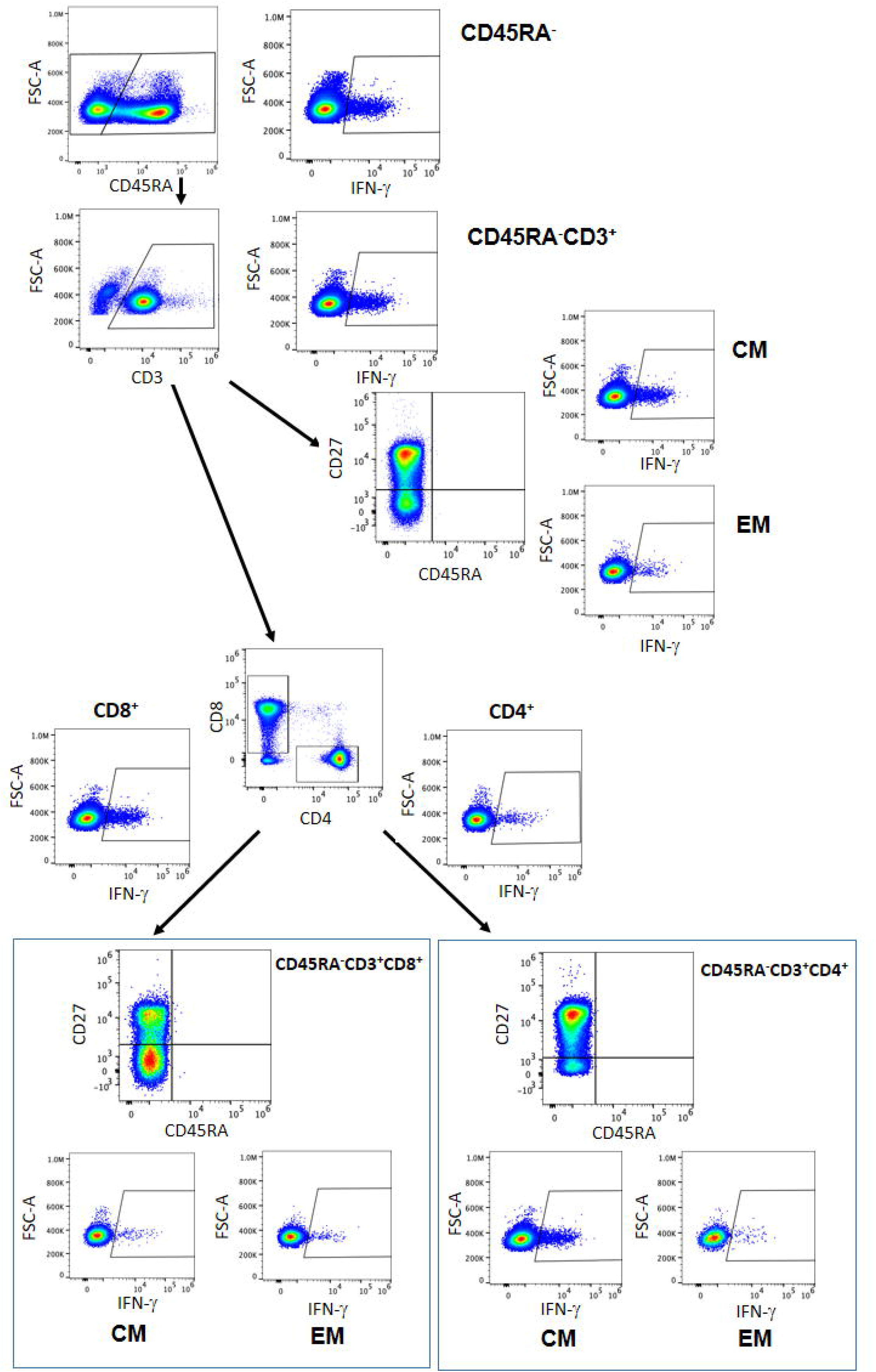
Representative figure of the gating strategy of the different cells subsets within the CD45RA^−^ population and the IFN-γ expressed. CM= central memory, EM=effector memory

### Large-scale CD45RA depletion. Creation of an off-the self-biobank of CD45RA-memory T cells from a COVID-19 convalescent donor

Blood donor apheresis was performed followed by CD45RA^+^ T cell depletion as described in Material and Methods. After depletion of CD45RA^+^ cells, 99.8% of the cells were CD45RA^−^CD3^+^. Cells were cryopreserved and stored. Most of the CD45RO^+^ cells were CD4^+^ with a high CD4/CD8 ratio. The viability after thawing was 98-99%.

### Identification of SARS-CoV-2 specific CD4^+^ T and CD8^+^ T cell responses within CD45RA^−^memory T cells after CD45RA depletion

After CD45RA depletion, the percentage of CD45RA^−^CD3^+^ cells was 80.05%. Within that population 83.56% of the cells were CD4^+^ and 14.37% were CD8^+^ T cells. After exposure to the three peptides specific for SARS-CoV-2, we found that CD3^+^, CD3^+^CD4^+^, and CD3^+^CD8^+^ subsets expressed IFN-γ with percentages of 0.36, 0.38, and 0.31 respectively. (**Table 3**).

**Table 3.** Phenotype of CD45RA memory T cells from a convalescent donor after large scale CD45RA depletion.

| | Cell marker | % of cells after thawing | % IFNγ<sup>+</sup> | Fold change IL15 (o/n)* | Fold change IL15 (72h)** |
| --- | --- | --- | --- | --- | --- |
| **CD45RA-subpopulations** | **CD45RA<sup>-</sup>** | 99.85 | 0.36 | 1.00 | 0.99 |
|  | CD45RA<sup>-</sup>CD3<sup>+</sup> | 80.05 | 0.36 | 1.05 | 0.99 |
|  | CD45RA<sup>-</sup>CD3<sup>+</sup>CD4<sup>+</sup> | 83.56 | 0.38 | 1.02 | 0.98 |
|  | CD45RA<sup>-</sup>CD3<sup>+</sup>CD8<sup>+</sup> | 14.37 | 0.31 | 0.84 | 1.10 |
|  | **CD45RA<sup>-</sup> CD4<sup>+</sup>** |  |  |  |  |
|  | CD27<sup>+</sup> (CM) | 89.13 | 0.38 | 1.01 | 1.02 |
|  | CD27<sup>-</sup> (EM) | 10.86 | 0.39 | 0.94 | 0.86 |
|  | CD127<sup>low</sup>CD25<sup>+</sup> (Treg) | 11.26 | N/A | 1.27 | 2.03 |
|  | **CD45RA<sup>-</sup> CD8<sup>+</sup>** |  |  |  |  |
|  | CD27<sup>+</sup> (CM) | 61.65 | 0.70 | 1.07 | 1.03 |
|  | CD27<sup>-</sup> (EM) | 38.34 | 0.00 | 0.89 | 0.95 |
| **T cell activation marker** | **CD3<sup>+</sup>** |  |  |  |  |
|  | HLA-DR<sup>+</sup> | 19.40 | 0.93 | 1.15 | 2.63 |
|  | CD69 BV421 high | 0.46 | 0.00 | 3.18 | 29.59 |
|  | CD25<sup>+</sup> | 60.68 | N/A | 1.50 | 8.52 |
|  | **CD4<sup>+</sup>** |  |  |  |  |
|  | HLA-DR<sup>+</sup> | 16.70 | 0.70 | 1.16 | 2.42 |
|  | CD69 BV421 high | 0.38 | 0.00 | 2.90 | 59.75 |
|  | CD25<sup>+</sup> | 66.10 | N/A | 1.48 | 8.58 |
|  | **CD8<sup>+</sup>** |  |  |  |  |
|  | HLA-DR<sup>+</sup> | 29.19 | 1.21 | 1.15 | 1.89 |
|  | CD69 BV421 high | 0.31 | 1.04 | 2.77 | 5.64 |
|  | CD25<sup>+</sup> | 9.60 | N/A | 4.42 | 36.42 |
| **T cell**<br>**Exhaustion**<br>**markers** | **CD3<sup>+</sup>** |  |  |  |  |
|  | NKG2A<sup>+</sup> | 1.88 | 0.00 | 0.73 | 1.39 |
|  | PD1 AF700 | 0.70 | 0.75 | 0.71 | 8.70 |
|  | **CD4<sup>+</sup>** |  |  |  |  |
|  | NKG2A<sup>+</sup> | 0.14 | 0.00 | 1.24 | 0.56 |
|  | PD1 AF700 | 1.43 | 0.62 | 0.84 | 9.15 |
|  | **CD8<sup>+</sup>** |  |  |  |  |
|  | NKG2A<sup>+</sup> | 10.24 | 0.00 | 0.76 | 1.22 |
|  | PD1 AF700 | 0.13 | 0.00 | 1.69 | 6.14 |
| **Chemokine**<br>**receptor and**<br>**integrin** | **CD3<sup>+</sup>** |  |  |  |  |
|  | CCR7 | 81.00 | N/A | 1.03 | 1.00 |
|  | CD103 | 2.18 | N/A | 0.74 | 1.24 |
|  | **CD4<sup>+</sup>** |  |  |  |  |
|  | CCR7 | 88.13 | N/A | 1.00 | 0.97 |
|  | CD103 | 1.38 | N/A | 0.74 | 0.99 |
|  | **CD8<sup>+</sup>** |  |  |  |  |
|  | CCR7 | 45.42 | N/A | 1.48 | 1.40 |
|  | CD103 | 5.96 | N/A | 0.89 | 1.39 |

\*Fold change increase after the treatment with IL15 overnight (o/n). \*\* Fold change increase after 72 hours with IL15. Fold change regarding the receptor expression has been calculated as the ratio between the value with IL15 and the value from a control without IL15. N/A= Not available

Within the EM and CM compartments, most of the CD4^+^ and CD8^+^ cells had a CM phenotype (89.1% v 61.6%). Both subpopulations CM and EM expressed IFN-γ after exposure with the three peptides. Within CD4^+^ CM, CD8^+^ CM and CD4^+^ EM subsets we found that 0.38%, 0.70% and 0.39% of cells respectively expressed IFN-γ. We didn’t find a specific expression of IFN-γ within the CD8^+^ EM subset. (Table 3).

Then, we characterized in-depth the phenotypic expression of activation, exhaustion, and Treg markers within the CD45RA^−^ population of the convalescent donor. The number of Treg defined by CD127^low^CD25^+^ was 11.26%. We found that the CD45RA^−^CD3^+^ expressed the activation marker HLA-DR (19.40%) in both subsets of cells CD4^+^ (16.70%) and CD8^+^ (29.19%). After exposure to the three SARS-CoV-2 specific peptides we found that within CD3^+^HLA^−^DR^+^, CD4^+^HLA^−^DR^+^, and CD8^+^HLA-DR^+^ populations 0.93%, 0.70% and 1.21% of the cells respectively were IFN-γ^+^. Also, the CD45RA^−^CD3^+^ expressed the exhaustion marker NKG2A (1.88%) and this expression was 10.24% within the CD8^+^ subset whereas it was nearly undetectable in CD4^+^ cells (0.14%). We didn’t find expression of IFN-γ within the NKG2A^+^ population. Also, 0.70% of the CD45RA^−^CD3^+^ cells expressed the exhaustion marker PD-1, this expression was 1.43% within the CD4^+^ subset and 0.13% within the CD8^+^ cells. We found 0.62% of IFN-γ^+^ cells within the CD45RA^−^CD3^+^CD4^+^PD-1^+^ population after exposure to the three peptides. (**Table 3**).

### CDR3 use of TCR

Polyclonal distribution for both TCR-β and -γ CDR3-encoding regions was almost identical among control and CD45RA^−^, whereas different oligoclonal fragments were seen in the CD45RA^+^ population. Three of the oligoclonal fragments seen in CD45RA^+^ cells were also identified in CD45RA^−^ (**Figure 4**).

**Figure 4.**
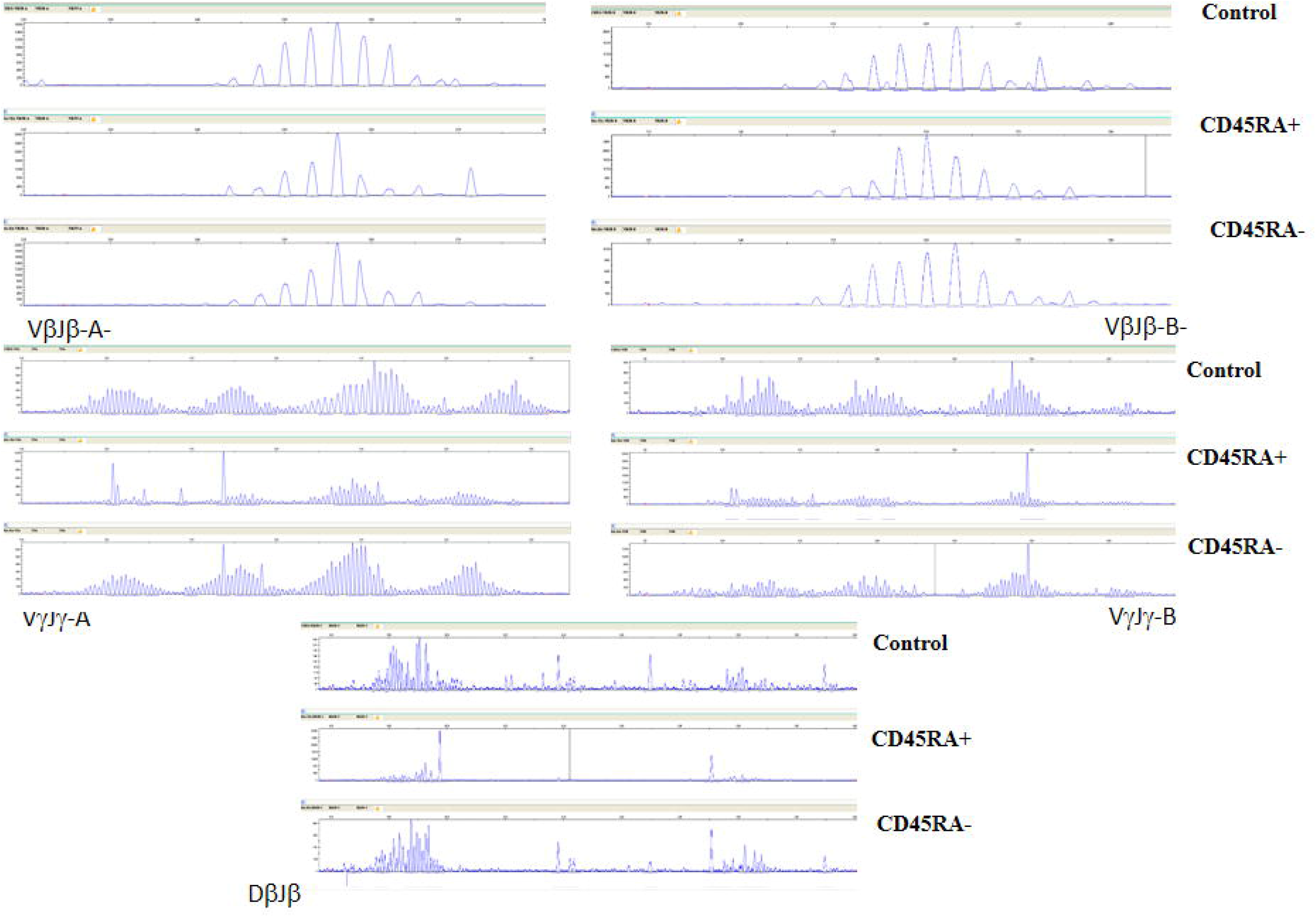
TCR-β and -γ spectratyping for 5 multi-primer sets covering most of the CDR3-encoding regions. Electropherograms showing gaussian distribution of polyclonal population (Control) is also mainly seen in CD45RA^−^, with oligoclonal patterns in CD45RA^+^. Some predominant fragments are shared in both convalescent donor populations.

### Induction of an activated memory T cells phenotype within CD45RA^−^ memory T cells after CD45RA depletion

IL15 is a cytokine fundamental for memory T cells that induces the activation, proliferation, and survival of T cells. To induce an activated phenotype of memory T cells we incubated the CD45RA-T cells with IL15 o/n and for 72 hours (**Table 3**). We observed a fold increase of the activation markers HLA-DR, CD69, and CD25 after 72 h of incubation (2.63, 29.59, 8.52) when compared to o/n incubation (1.15, 3.18, 1.50). The expression of the exhaustion markers NKG2A and PD-1 were also higher after 72 hours of incubation (**Supplementary Figures 1-5**). Then, we looked at the expression of chemokine CCR7 and the integrin CD103 important for homing to the air tract respiratory^26,27^. We observed that most of the CD3^+^CD4^+^ cells expressed CCR7^+^ cells (88.13%) and the CD3^+^CD8^+^ subpopulation expressed 45.42% of CCR7^+^ cells. The expression of CD103 was low in peripheral blood as expected^26^. We detected that the expression in the CD3^+^CD103^+^, CD3^+^CD8^+^CD103^+^ and CD3^+^CD4^+^CD103^+^ compartments was 2.18%, 5.96% and 1.38% respectively. Although the fold increase was not very remarkable in the CD45RA^−^CD3 ^+^ cells after 72 hours of incubation (1.00-fold CCR7 and 1.24-fold CD103), the increase was higher (1.40) within the CD8^+^ subpopulation (**Table 3 and Supplementary Figures 6, 7**).

## DISCUSSION

In the absence of an effective vaccine, and with the emergence of a second wave there is an urgent need to find effective treatments for COVID-19 disease. Here we report the existence of a SARS-CoV-2 specific T cell population within the CD45RA^−^ T memory cells of the blood from the convalescent donors. These cells can be easily, effectively, and rapidly isolated following a donor selection strategy based on the expression of IFN-γ after exposure with SARS-CoV-2 specific peptides and HLA antigen expression. This way we can obtain clinical-grade CD45RA^−^ memory T lymphocytes from the blood of convalescent donors, biobanking, thawing, and using them as a treatment for moderate/severe cases of COVID-19. These so-called “living drugs” retain memory against SARS-CoV-2 and other pathogens they have encountered previously. Unlike plasma where the concentration decreases after infusion, memory T cells expand and proliferate, so we hypothesize that will have a more lasting effect.

In previous studies this population of CD45RA^−^ memory T cells have shown no alloreactivity when compared to the CD45RA^+^ counterpart^28^, they are mainly CD4^+^29 and they have shown to be effective against viral infections^20^. Phenotypically we found that CD45RA^−^ memory T cells were fully capable of producing IFN-γ in the presence of SARS-CoV-2 specific peptides. Both CD4^+^ and CD8^+^ CM and EM subsets were able to generate IFN-γ after exposure to the SARS-CoV-2 peptides showing coverage of response. CD8^+^ cytotoxic cells can kill virally infected cells by secreting cytokines, at the same time CD4^+^ T cells increase the ability of the CD8^+^ T cells in eliminating the virus. They have been shown to play an important role in controlling the viral replication of other viruses such as EBV and CMV^30^. EM are the first responders to the infection with a quick and strong response to pathogens whereas CM proliferate and create a new round of effector T cells^22,31^. IL15 is essential for the survival of memory CD8^+^ and CD4^+^ T cells subsets promoting the activation of CD4^+^T cells including cytokine production, proliferation, and maintenance of memory population^32,33^. After incubating for 3 days with IL15 we obtained phenotype characteristic of an activated state as is shown by the fold increase in the activation markers HLA-DR, CD69, and CD25 and in CCR7 and CD103 markers characteristic of homing to the lymph nodes and mucosal tissues.

In our study, we did not observe the production of IFN-γ by SARS-Cov-2 specific T cells in healthy unexposed individuals which is in agreement with the data from Peng Y et al^34^ but differs from previously published data^14^. This could also due to the different detection methods used and the small sample size.

Studies have shown that the correlation between neutralizing antibodies and the severity of the symptoms where antibody responses wane over time even in a short period as 6-7 weeks after the onset of symptoms^35–37^. Importantly our data shows the existence of the SARS-CoV-2 memory T cells in convalescent donors with mild symptoms. This has a great implication for the protection from the next SARS-CoV-2 infections and to decrease the severity of COVID-19 disease. Further studies with a larger cohort to determine the duration of memory to SARS-CoV-2 will be needed to elucidate long-term protection to SARS-CoV-2 as it has been shown for another coronavirus previously^16^.

For the proper T cell recognition, both donor and recipient need to share HLA alleles. Given the vast number of convalescent donors to find a right haploidentical donor based on HLA typing would not be difficult. We estimate that based on HLA donor-recipient match four donors will cover almost the whole country population^38^.

The procedure to obtain the cells is easy to implement for small scale manufacture, quick and cost-effective involving minimal manipulation and without GMP condition requirements. These factors make feasible the generation of a biobank or stock from the blood of convalescent donors that would be immediately available ‘off the shelf’ for subsequent outbreaks of the pandemic increasing the therapeutic options available in the current SARS-CoV-2 pandemic. The safety of these cells in moderate/severe patients with COVID-19 is currently assessed in a clinical trial (NCT04578210).

These cells could provide to the patient: 1) a pool of SARS-CoV-2 specific T cells that will respond quickly to the infection, 2) a pool of lymphocytes to patients with severe disease presenting with lymphopenia and 3) a pool of specific memory T cells for other pathogens from the donors encountered in their life which are vital to clear other secondary infections usually developed in hospitalized patient COVID-19^39,40^.

## Contributors

CF, BS, and A P-M designed the study. CF, BP-M and CM-D performed the in vitro experiments. JLV, AB, FGS performed HLA typing and TCR spectratyping. R D-Paz and AM performed the non-mobilized apheresis and cryopreservation of the cells. CM performed the statistics. CF, BP-M, BS and A P-M wrote the first draft of the manuscript. All authors revised the manuscript, participated in the interpretation of the data, approval of the manuscript, and submission.

## Acknowledgments

We thank the donors for their blood donation and the CRIS Cancer Foundation (http://criscancer.org) and the Generalitat Valenciana AVI-GVA COVID-19-68 for their support.

## Conflict of interest

CF, BS and AP-M filed a patent on this topic.

BS received fees from Celgene, Gilead, Sanofi and Novo-Nordisk not related to this work.

---

**Supplemental Figure 1**. Expression of HLD-DR in the CD45RA^−^CD3^+^, CD45RA^−^ CD3^+^CD4^+^ and CD45RA^−^CD3^+^CD8^+^ populations from the convalescent donor. Cells were culture with 50 ng/ml of IL15 o/n and for 72 hours.

**Supplemental Figure 2**. Expression of CD69 in the CD45RA^−^CD3^+^, CD45RA^−^CD3 ^+^CD4^+^ and CD45RA^−^CD3^+^CD8^+^ populations from the convalescent donor. Cells were culture with 50 ng/ml of IL15 o/n and for 72 hours.

**Supplemental Figure 3**. Expression of CD25 in the CD45RA^−^CD3^+^, CD45RA^−^CD3 ^+^CD4^+^ and CD45RA^−^CD3^+^CD8^+^ populations from the convalescent donor. Cells were culture with 50 ng/ml of IL15 o/n and for 72 hours.

**Supplemental Figure 4**. Expression of NKG2A in the CD45RA^−^CD3^+^, CD45RA^−^CD3 ^+^CD4^+^ and CD45RA^−^CD3^+^CD8^+^ populations from the convalescent donor. Cells were culture with 50 ng/ml of IL15 o/n and for 72 hours.

**Supplemental Figure 5**. Expression of PD-1 in the CD45RA^−^CD3^+^, CD45RA^−^ CD3^+^CD4^+^ and CD45RA^−^CD3^+^CD8^+^ populations from the convalescent donor. Cells were culture with 50 ng/ml of IL15 o/n and for 72 hours.

**Supplemental Figure 6**. Expression of CCR7 in the CD45RA^−^CD3^+^, CD45RA^−^CD3^+^CD4^+^ and CD45RA^−^CD3^+^CD8^+^ populations from the convalescent donor. Cells were culture with 50 ng/ml of IL15 o/n and for 72 hours.

**Supplemental Figure 7**. Expression of CD103 in the CD45RA^−^CD3^+^, CD45RA^−^ CD3^+^CD4^+^ and CD45RA^−^CD3^+^CD8^+^ populations from the convalescent donor. Cells were culture with 50 ng/ml of IL15 o/n and for 72 hours.

