## Supplementary figures and images for "SARS-CoV-2 specific memory T lymphocytes from COVID-19 convalescent donors: identification, biobanking and large-scale production for Adoptive Cell Therapy"

### Supplemental Figure 1

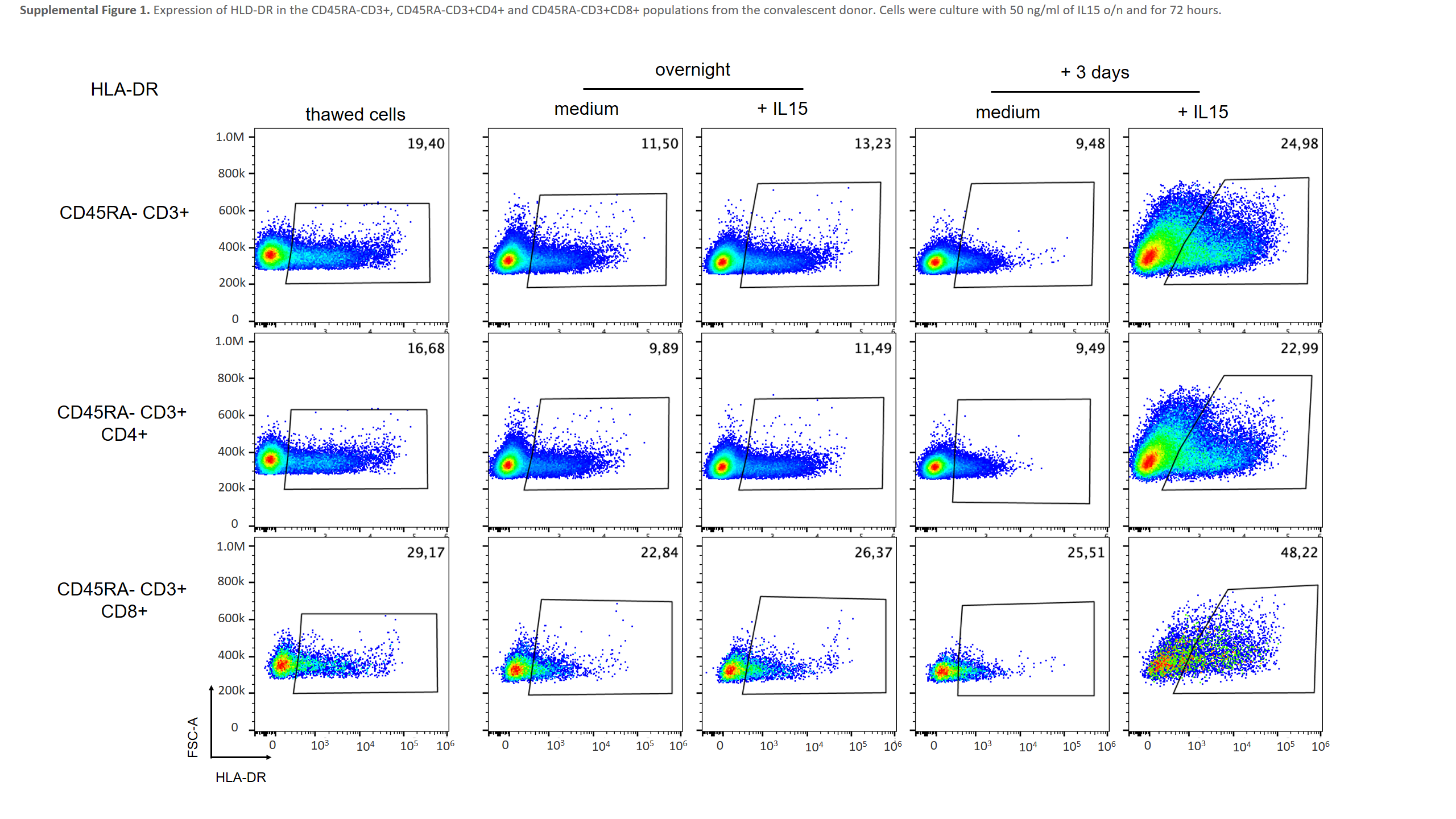

### Supplemental Figure 2

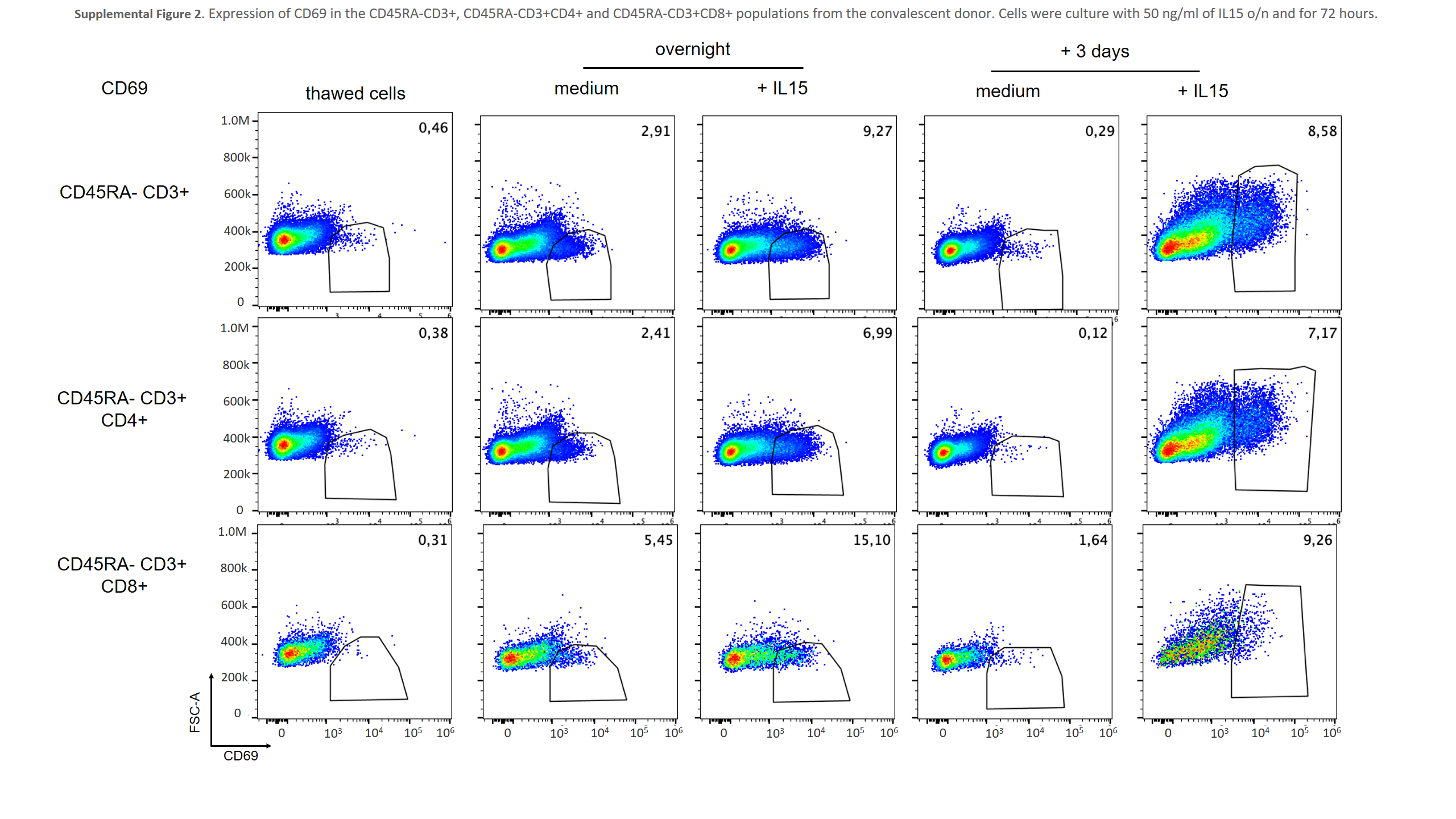

### Supplemental Figure 3

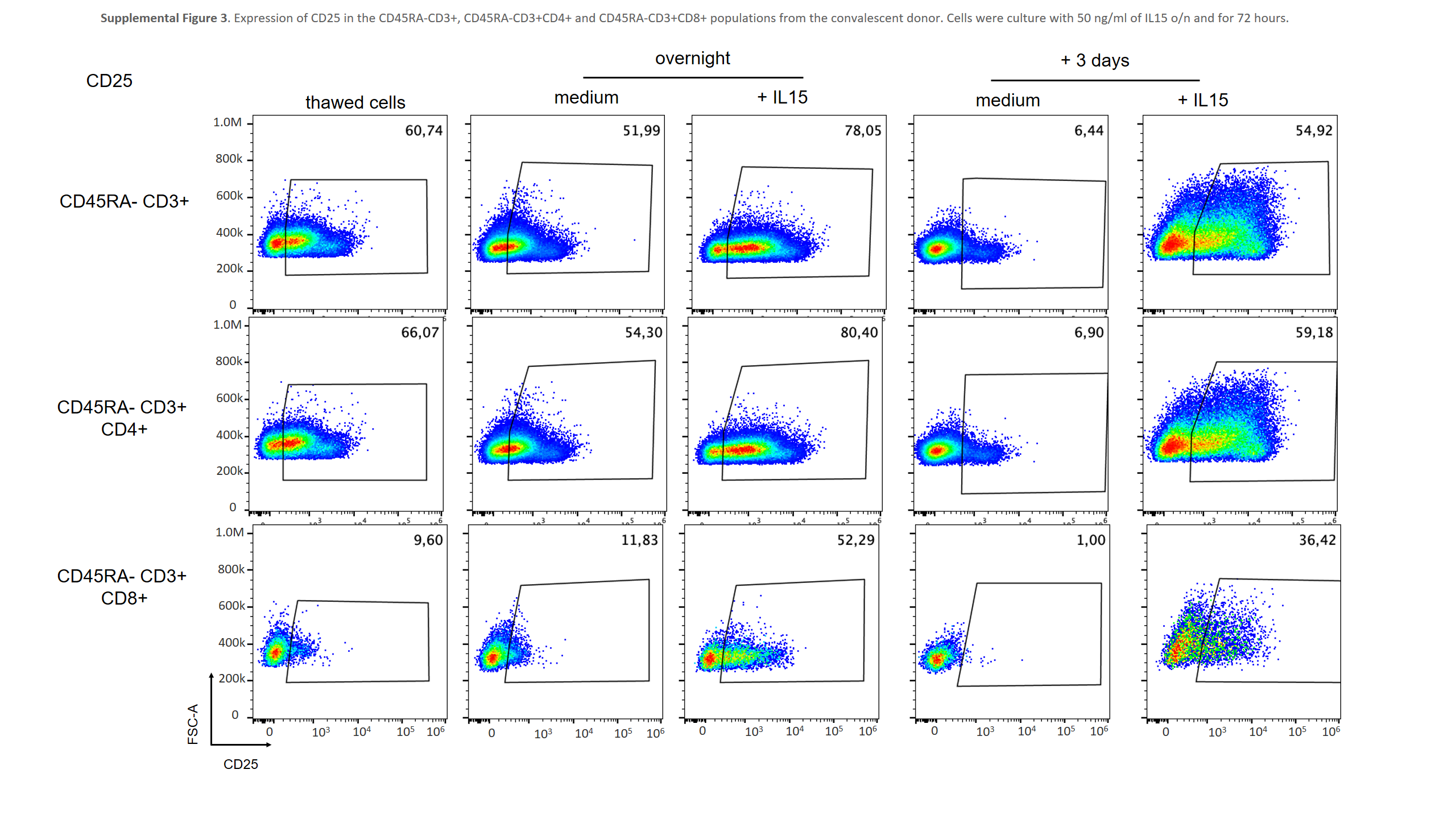

### Supplemental Figure 4

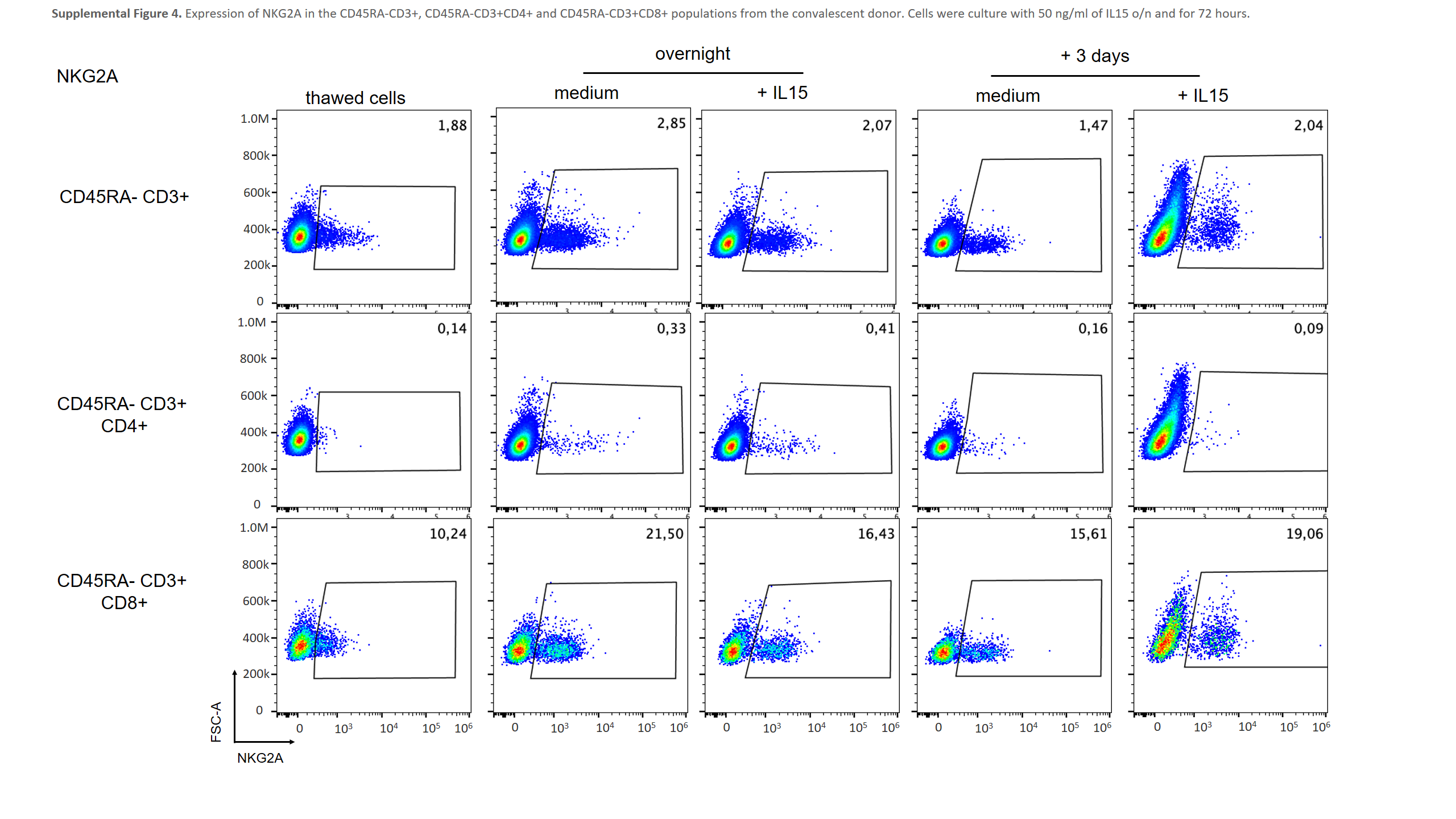

### Supplemental Figure 5

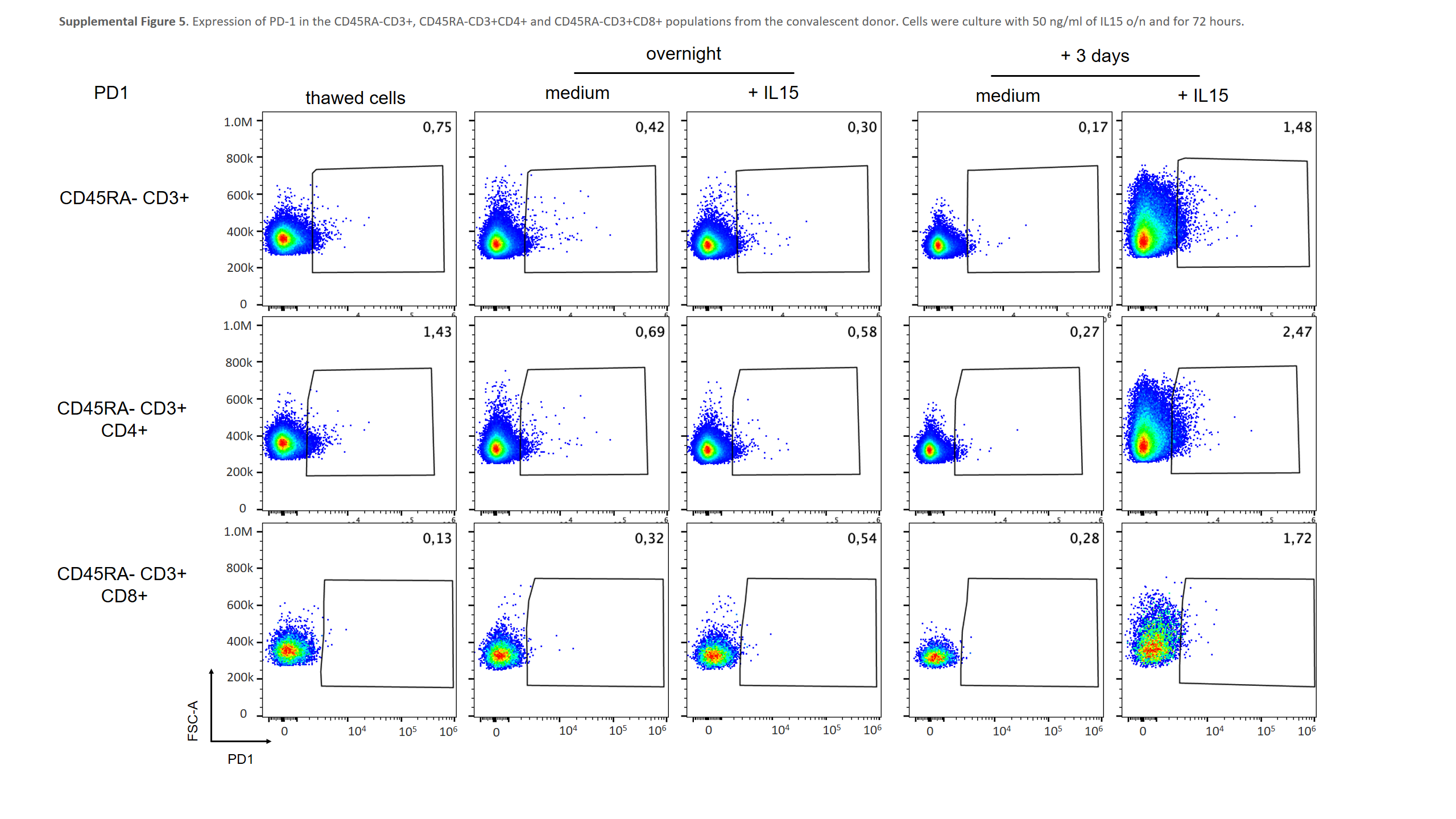

### Supplemental Figure 6

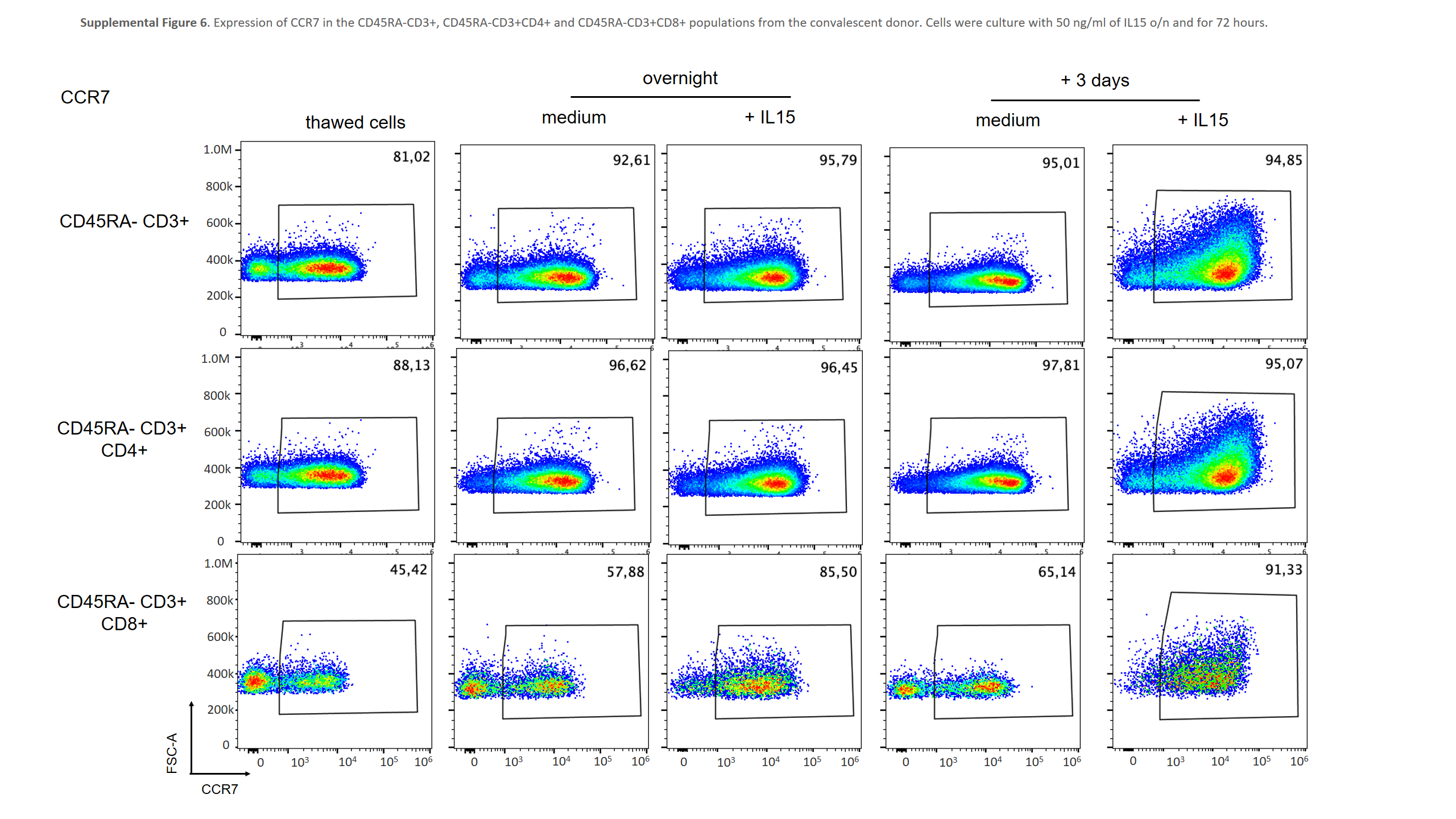

### Supplemental Figure 7

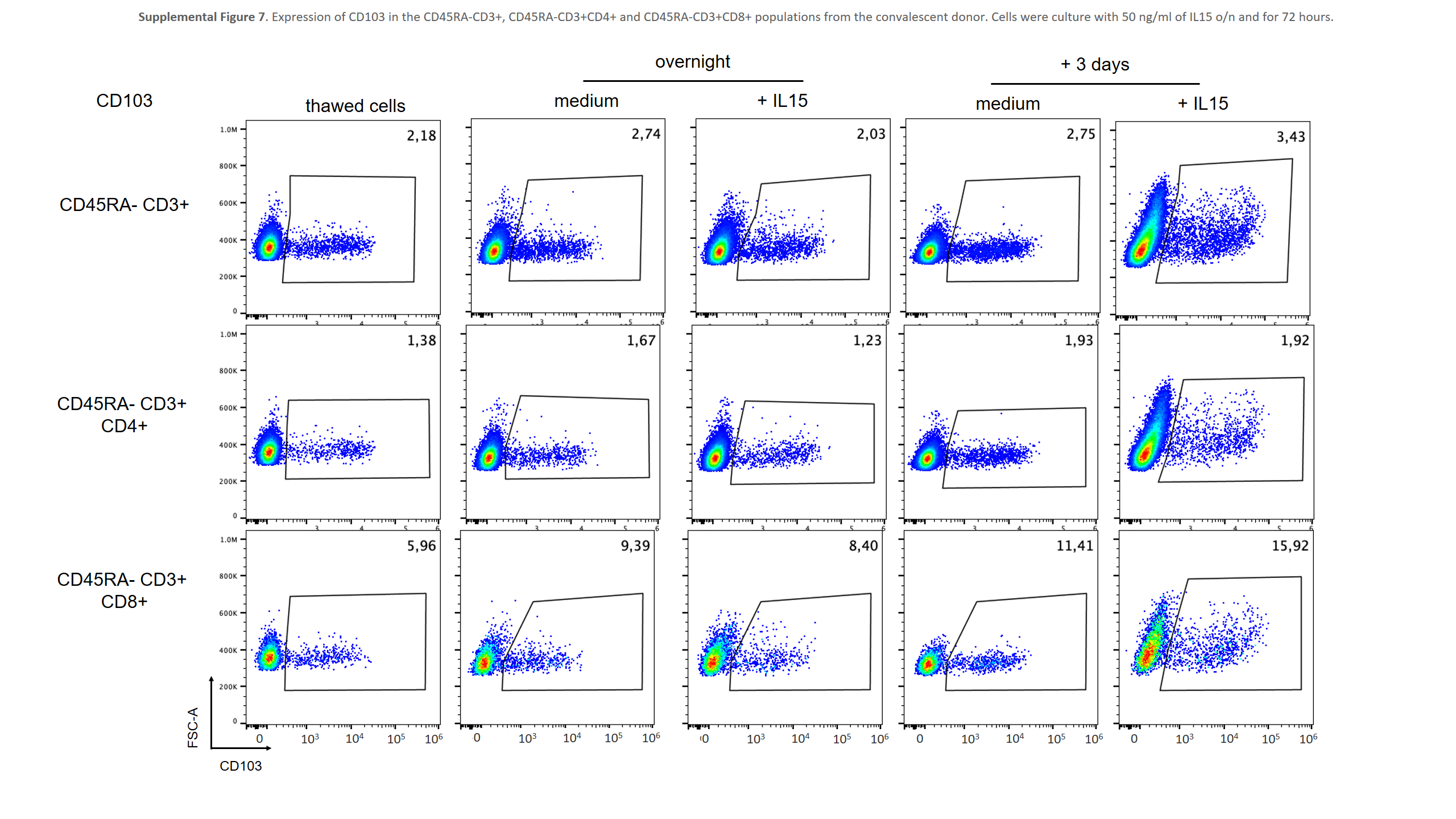
